# Nucleus accumbens core single cell ensembles bidirectionally respond to experienced versus observed aversive events

**DOI:** 10.1101/2023.07.17.549364

**Authors:** Oyku Dinckol, Jennifer E. Zachry, Munir Gunes Kutlu

## Abstract

Empathy is the ability to adopt others’ sensory and emotional states and is an evolutionarily conserved trait among mammals. In rodents, empathy manifests itself as social modulation of aversive stimuli such as acknowledging and acting on conspecifics’ distress. The neuronal network underlying social transmission of information is known to overlap with the brain regions that mediate behavioral responses to aversive and rewarding stimuli. In this study, we recorded single cell activity patterns of nucleus accumbens (NAc) core neurons using in vivo optical imaging of calcium transients via miniature scopes. This cutting-edge imaging methodology not only allows us to record activity patterns of individual neurons but also lets us longitudinally follow these individual neurons across time and different behavioral states. Using this approach, we identified NAc core single cell ensembles that respond to experienced and/or observed aversive stimuli. Our results showed that experienced and observed aversive stimuli evoke NAc core ensemble activity that is largely positive, with a smaller subset of negative responses. The size of the NAc single cell ensemble response was greater for experienced aversive stimuli compared to observed aversive events. Our results also revealed a subpopulation within the NAc core single cell ensembles that show a bidirectional response to experienced aversive stimuli versus observed aversive stimuli (i.e., negative response to experienced and positive response to observed). These results suggest that the NAc plays a role in differentiating somatosensory experience from social observation of aversion at a single cell level. This has important implications for psychopathologies where social information processing is maladaptive, such as autism spectrum disorders.

## Introduction

Empathy is the ability to adopt others’ sensory and emotional state and can be demonstrated by diverse behaviors such as mimicry, emotional contagion, and altruistic behavior ^1, 2^. Empathy is an evolutionarily conserved trait among mammals and the phenotypes related to empathy have been shown in rodents (i.e., observational fear, social modulations of pain^34^). Observational fear is a well-known and widely utilized paradigm for assessing empathy-like behavioral phenotypes, which can be characterized as forming an association between an aversive stimulus and an outcome via social means.

Perception of social information may be maladaptive in certain psychopathologies. Studies have shown altered neuronal activities in overlapping brain-wide networks during social stimuli in psychiatric pathologies such as autism spectrum disorder (ASD) and schizophrenia^5–7^. In addition, forming an association between a neutral stimulus and an aversive outcome through social means in the absence of direct experience likely contributes to the etiology of anxiety disorders^8^. Anxiety disorders are common companions of neurodevelopmental conditions such as ASD^9^ as one of the leading factors of anxiety in individuals with ASD. Thus, in order to further our understanding of these pathologies, it is important to explore the neural underpinnings of aversive learning through social means.

The neuronal network underlying social information processing is known to overlap with the brain regions that mediate the aversive stimuli response and reward circuitry^10, 11^. Dopaminergic and GABAergic circuits have been shown to underlie the neural bases of observational aversive learning including brain regions such as the anterior cingulate cortex (ACC), the amygdala, and the nucleus accumbens (NAc)^12–15^. The social transfer of fear has been linked to NAc projections from the ACC^1, 16^. The NAc is a heterogeneous, limbic structure that relates to associative learning and conditioned behavioral responses; therefore, it plays a critical role in behavioral control^17^. Despite serving as a critical node in the brain’s reward circuitry, the NAc is also critically involved in modulating responses to aversive stimuli^17, 18^. The NAc comprises subregions, the core and the shell, that are known to play dissociable roles^17, 19–24^. Studies have shown that the NAc core is one of the main mediators of aversive learning^19^. It has also been previously shown that the NAc is involved in the acquisition of fear responses^17^ however the involvement of the NAc specifically on observational aversive learning remains mostly underexplored.

In this paper, we aimed to examine NAc core single cell ensemble activity in response to experiencing versus observing aversive events by performing *in vivo* optical imaging of calcium activity via miniature scopes. This approach is uniquely suited for this aim because, unlike electrophysiological methods, it allows us to measure the activity from single cells and longitudinally follow each cell and its activity under different behavioral conditions. Using this approach, we tested mice in two separate behavioral contexts where they first experienced a set of footshocks followed by a session where they observed a conspecific receiving the same footshocks. In this way we aimed to examine NAc core single cell activity when animals experience versus observed aversive stimuli.

Our results revealed a larger NAc core response to experienced shocks compared to observed shocks. In the experienced shock condition, there was a large single cell ensemble of positively responding cells and this was reduced to a smaller ensemble in the observed shock condition. These single cell calcium responses did not reveal a habituation pattern throughout the shock presentations as these responses remained relatively stable throughout the footshock presentations. However, our longitudinal analysis of single cells revealed that the NAc core cells that respond negatively to the experienced footshocks switched to responding positively when those shocks were observed. In sum, our results suggest that a sub-population in the NAc core single cell ensembles exhibit a bidirectional response pattern to experienced versus observed aversive stimuli.

## Results

### Experienced aversive stimuli evoke a larger NAc core ensemble activity compared to observed aversive events

In this study, we aimed to identify single cell ensembles within the NAc that respond to experienced and/or observed aversive events. A group of C57BL/6J mice (observer mice) were subjected to a set of footshocks in an operant box and the NAc core single cell activity was recorded by using *in vivo* optical imaging of calcium activity via miniature scopes during the “experienced shock session” (see **Supp. Fig. 1** for GRIN lens placements used for the endoscopic miniature scopes and see **Supp. Fig. 2** for a representative cell map and cell traces). Twenty-four hours later, during the “observed shock session”, cell responses from the same animals were recorded while they were observing another same-sex partner (performer mice) experience the footshocks (**Fig.1 a**). During each session, animals were subjected to 17 footshocks with the same inter-stimulus interval (**Fig.1 b**).

**Figure 1.**
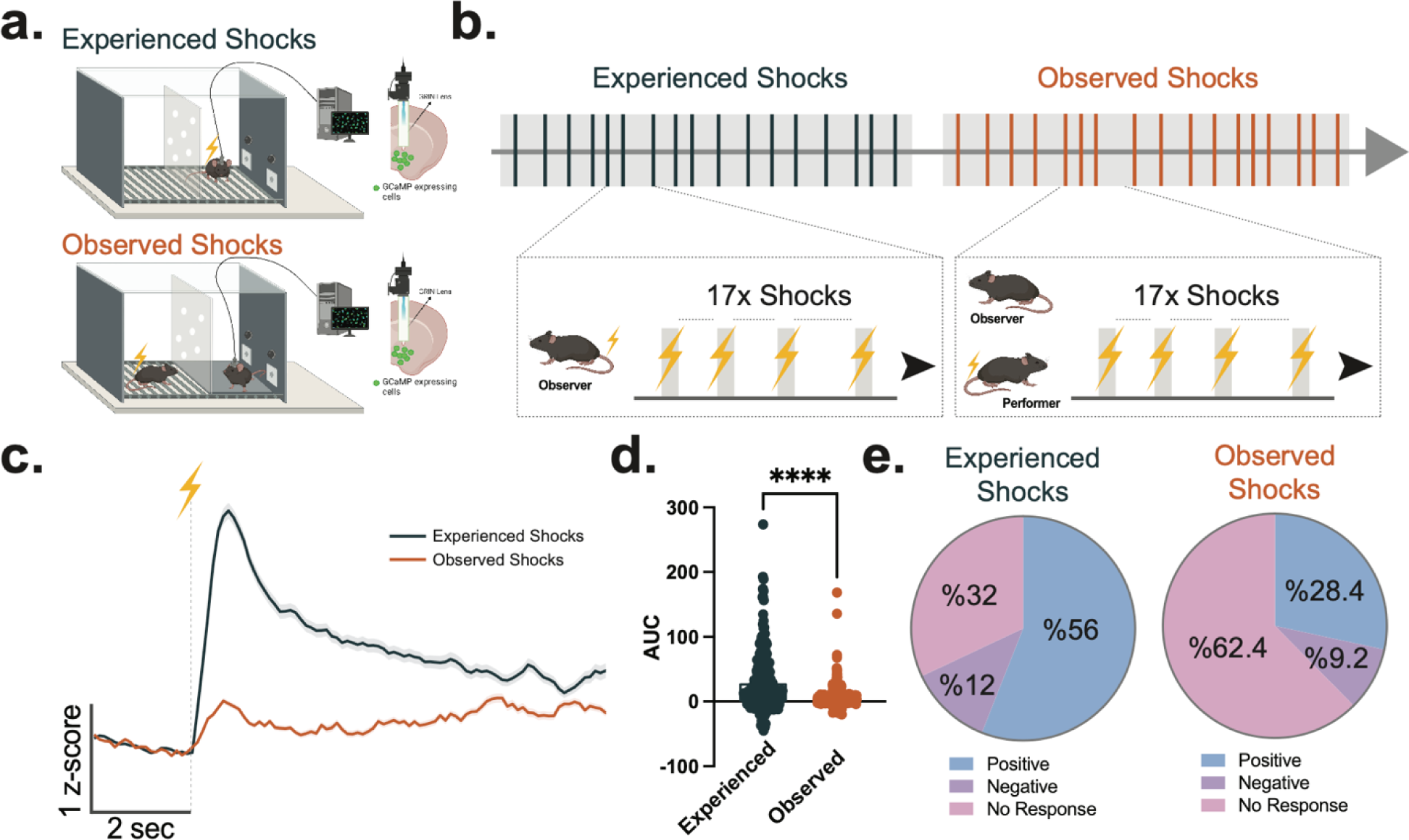
NAc core single cell ensembles respond to both experienced and observed aversive events. **(a)** NAc core single cell activity was recorded via miniscopes combined with a calcium sensor (GcaMP6m) while mice received or observed another mouse receiving aversive footshocks. **(b)** Schematic demonstration of the footshock presentations during the experiment. In each session, mice either received 17 footshocks or observed another mouse receiving 17 footshocks. **(c)** Mean cell response to the experienced versus observed footshock presentations across all cells from all animals. **(d)** Mean area under the curve (AUC) for the population response was larger to the experienced compared to observed footshock events (unpaired t-test, *t*_544_=9.585, *p*<0.0001, n=250-396 cells; independent t-tests against the theoretical mean=0 AUC for experienced footshocks *t*_249_=10.43, *p*<0.0001, mean=29.61; for observed footshocks *t*_395_=7.618, *p*<0.0001, mean=6.002). **(e)** The experienced and observed footshock responses show bidirectionality. (**left**) Percentages of the cells showing a positive (56%), negative (12%), or no response (32%) to the experienced footshock. (**right**) Percentages of the cells showing a positive (28.4%), negative (9.2%), or no response (62.4%) to the observed footshock. Data represented as mean ± S.E.M. **** *p* < 0.0001.

Our results revealed that both experienced and observed footshock elicited a positive population response (**Fig.1 c,d**). However, the experienced aversive stimuli evoked a significantly greater NAc core single cell ensemble response compared to the observed aversive events (**Fig.1 c,d**). The experienced aversive stimuli elicited a large ensemble of positive cell activity (56% of all detected cells); meanwhile there was a smaller portion of negative (12%) or no response (32%) cell ensembles. On the other hand, the observed aversive events also evoked a positive NAc core single cell ensemble response, but this response was relatively weaker, and the size of the ensemble was relatively smaller compared to the experienced aversive stimuli ensemble (28.4%). However, the size of the negative responses was relatively similar in both events (12% for experienced versus 9.2% for observed footshocks; **Fig.1 e**). Moreover, a separate comparison of the positive and negative responder cell activity revealed that the experienced aversive stimuli evoked larger positive and negative responses compared to the observed aversive events (**Supp. Fig. 3**).

We also wanted to examine if these cell responses habituate with repeated presentations of the experienced or observed aversive stimuli. The NAc core single cell responses showed a relatively stable pattern for each footshock presentation. Each experienced aversive stimulus evoked a similar trend of NAc core activity throughout the behavioral session. Likewise, each observed aversive event evoked a similar NAc core response. Therefore, the ensemble response to the experienced and observed aversive events was not habituated and remained relatively stable (**Fig. 2 a,b**). These results demonstrate that NAc core single cell ensembles not only respond to experienced aversive events but also respond to aversive events that are observed, suggesting that aversive information encoding within the NAc core has a social dimension.

**Figure 2.**
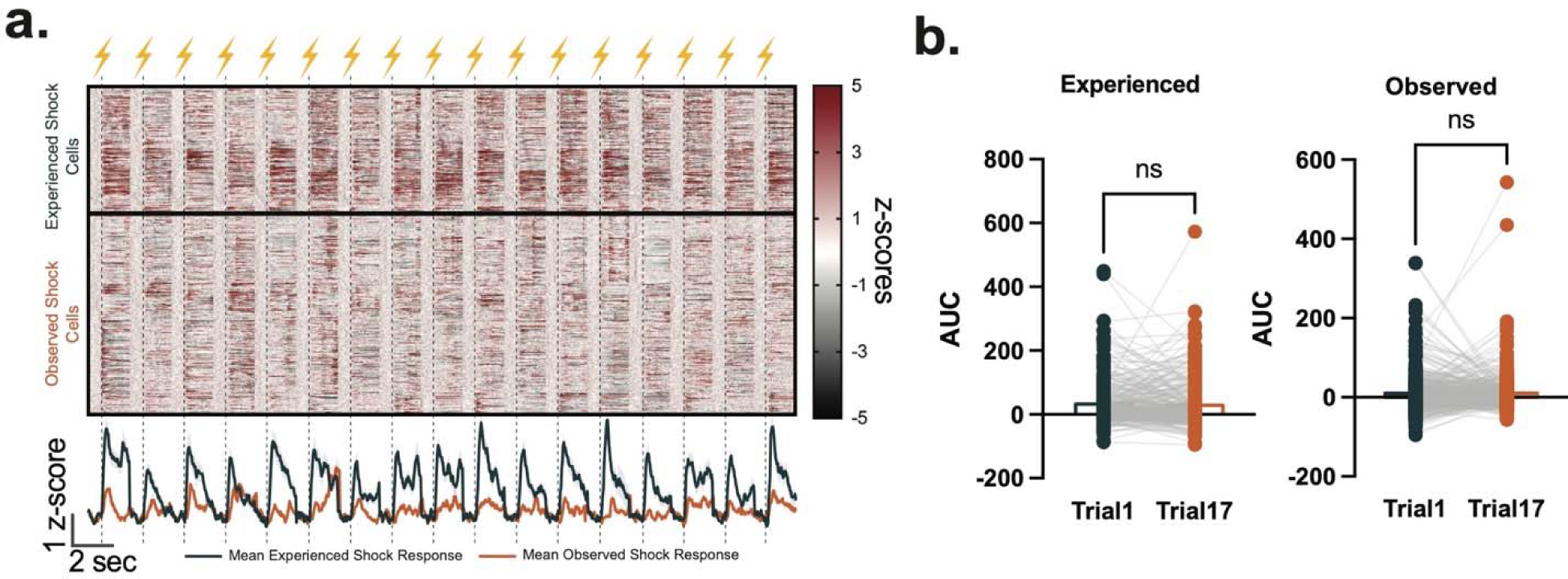
Single cell responses to the experienced and observed footshocks did not habituate with experience. (**a**) Heatmap showing single cell responses from individual cells (**upper panel**; z-scores) and mean single cell responses (**lower panel**; z-scores) for each aversive event throughout each session. (**b**) The size of the single cell responses (Area under the curve, AUC) did not differ between the first and last (17^th^) experienced (**left panel**; paired t-test, *t*_249_=0.6081, *p*=0.5437, n=250 cells) or observed (**right panel**; paired t-test, *t*_395_=0.1279, *p*=0.8983, n=396 cells) footshock. Data represented as mean ± S.E.M. ns = *p* > 0.05.

### A subpopulation within the NAc core single cell ensembles show a bidirectional response to experienced versus observed aversive events

Next, we wanted to follow up each cell longitudinally in order to examine their specific responses to the events where the animal experienced aversive stimuli versus observed aversive stimuli. Longitudinal registration analysis revealed that a large number of NAc core cells were active during both experienced and observed aversive events. About 50% of the cells detected during the experienced shock session (127 out of the 250 cells detected) and 32% of the cells detected during the observed shock session (127 out of the 396 cells detected) were co-registered (**Supp. Fig. 4**). Importantly, a subpopulation in the NAc core single cell ensembles exhibited a bidirectional response pattern to experienced versus observed aversive stimuli. Specifically, a number of NAc core single cells that show a positive response to experienced footshock switched to showing the opposite response when the animals observed the same aversive event (**Fig. 3 a,b**). A majority of the NAc core single cells (73%; **Supp. Fig. 5**) that showed a negative response to experienced footshock showed a positive response when the animals observed the same aversive event (**Fig. 3 a,b**). We found a similar bidirectional response trend for the cell ensembles that show a negative response to observed aversive events (**Fig. 3 c,d**). However, the cells that exhibited a positive response to observed aversive events tended to show a similar positive response to experienced aversive stimuli (**Fig. 3 c,d**).

**Figure 3.**
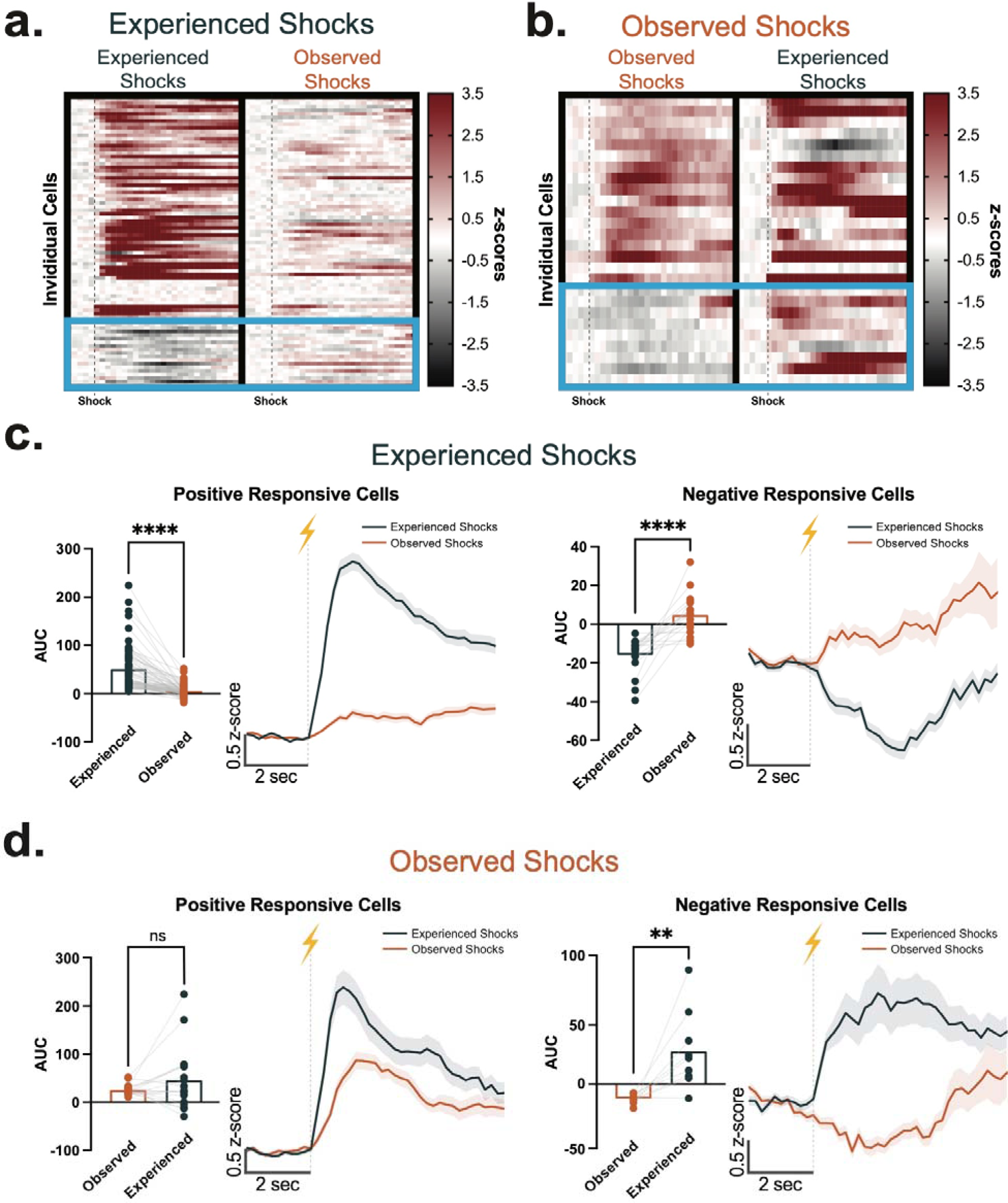
A sub-population of the NAc core single cell ensembles show a bidirectional response pattern to experienced versus observed aversive events. Heatmaps showing responses to (**a**) experienced versus (**b**) observed footshocks from the NAc core single cell sub-population with significant positive or negative AUCs. The blue boxes highlight the cells that exhibited bidirectional response, which was a switch from a negative to positive response. (**c**) Averaged cell responses depicted as mean AUCs (**left**) and averaged z-scores (**right**) from the cells that showed significant positive or negative responses to the *experienced* footshocks. The cells that showed a significant *positive* response to the experienced footshocks showed significantly weaker responses to the observed footshocks (paired t-test, *t*_62_=8.431, *p*<0.0001, n=63 cells). The cells that showed a significant *negative* response to the experienced footshocks showed significantly stronger responses for the observed footshocks (paired t-test, *t*_16_=5.298, *p*<0.0001, n=17 cells). (**d**) Averaged cell responses depicted as mean AUCs (**left**) and averaged z-scores (**right**) from the cells that showed significant positive or negative responses to the *observed* footshocks. The cells that showed a significant *positive* response to the observed footshocks showed similar size responses to the experienced footshocks (paired t-test, *t*_16_=1.289, *p*=0.215, n=17 cells). The cells that showed a significant *negative* response to the observed footshocks showed significantly stronger responses for the experienced footshocks (paired t-test, *t*_8_=3.441, *p*<0.0088, n=9 cells). Data represented as mean ± S.E.M. ** *p* < 0.01, **** *p* < 0.0001.

For an unbiased examination, we analyzed the longitudinal cell activities via hierarchical clustering. Our analysis revealed comparable results, which also showed that there were subgroups of NAc core cells that showed a bidirectional response to the experienced versus observed aversive stimuli (**Fig. 4**). These results suggest that there are specific subgroups of NAc core single cell ensembles that bidirectionally encode experienced versus observed aversive events. Thus, it is possible that NAc core single cells are functionally divergent depending on the source of the aversive information (e.g., social versus somatosensory).

**Figure 4.**
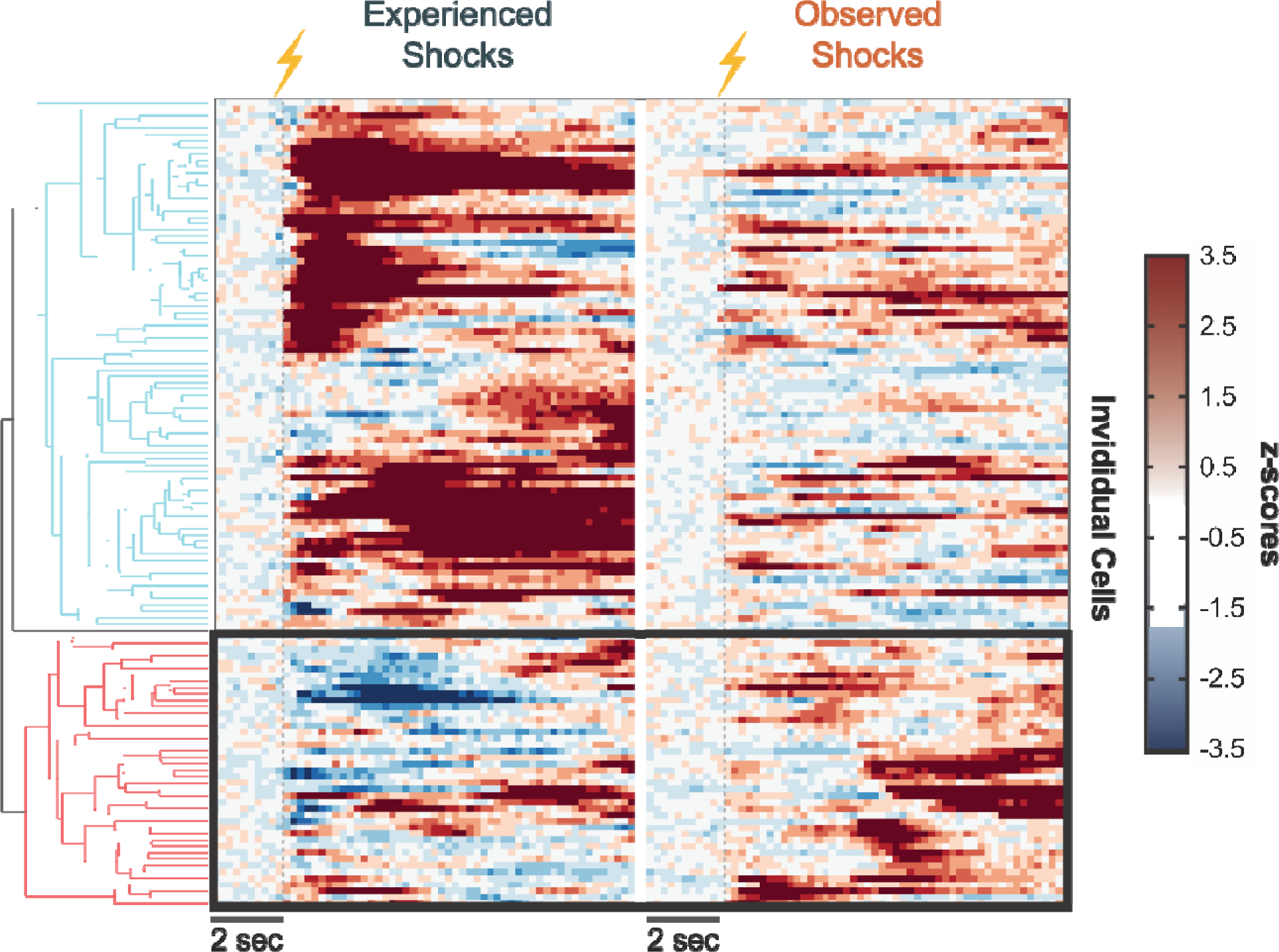
Unsupervised clustering algorithms group cells based on bi-directional response patterns. **(a)** Hierarchical clustering identified two major clusters of NAc core single cells based on their calcium responses: 1) Cells that show positive responses (shades of red) to the experienced footshock versus 2) Cells that show negative responses (shades of blue) to the experienced footshock. The second cluster mostly switched their response pattern to a positive response when the mice observed the same aversive event.

## Discussion

In the present study, we aimed to identify single cell ensembles within the NAc core that respond to aversive stimuli via using cutting-edge approaches of *in vivo* optical imaging of calcium activity via miniature scopes. This methodology offers unique advantages for single neuron recordings as we were able to follow the activity patterns of NAc single cells throughout different behavioral sessions and experiences. Taking advantage of this methodology, we compared the activity patterns of NAc single cells when the mice experienced aversive stimuli versus observed aversive stimuli. Our results showed that experienced aversive stimuli evoke a NAc core ensemble activity that is largely positive with a smaller portion of negative responses. Interestingly, observed aversive events also elicited a significant positive response, however this response was weaker. The size of the positively responding single cell ensembles was greater for experienced aversive stimuli compared to observed aversive events. These results are in line with the previous studies showing that the NAc mediates aversive stimuli through evoking neuronal activities, altered neuromodulation and gene expression, as well as behavioral responses^25–38^. Our results similarly showed that most NAc core single cells positively respond to aversive stimuli, however, there was also a smaller group of cells exhibiting a negative response to experienced aversive stimuli suggesting that the NAc core single cell ensembles are heterogeneous in their response to aversive stimuli.

Importantly, we found that not only experiencing aversive stimuli, but also merely observing other mice experiencing aversive stimuli evokes NAc core single cell responses time-locked to the aversive event. This suggests that aversive information obtained through both somatosensory and social perception are processed within the NAc single cell ensembles. A number of studies have shown that aversive information such as fear can be learned through observation and the neural basis of observational fear consists of the overlapping circuitry of social behavior and reward/aversive responses^1, 11–13, 17^ including processing within the NAc^3, 10, 14, 39^. Studies in mice showed altered NAc activity in the course of social interactions^10, 40^. Similarly, a functional magnetic resonance imaging study in humans showed NAc recruitment on social motivation (i.e., approval, disapproval^39^). Our results are in line with these previous reports, however they show that the NAc core single cell responses are less homogenous than it was previously thought.

In addition to confirming that the NAc core is heavily involved in the processing of experienced and observed aversive stimuli, we critically found that there is a sub-population of NAc single cells that shows bidirectional response patterns to experienced versus observed aversive events. That is, a group of NAc core cells negatively respond to experienced and positively respond to observed aversive events (or vice versa). There are multiple ways we can interpret this bidirectional computation within the NAc core. First, it is possible that this computation may be a part of the neural representation of self-perception where the organism differentiates between events happening to “self” versus “others”. However, based on previous literature on the perception of “self”^41, 42^ it is likely that self-perception requires a widespread coordinated activity involving a larger network within the brain. Therefore, it is unlikely that the pattern of activation we observed in the NAc single cells is unique to this brain region and it may be observed in other brain regions. It is also possible that the differential activity patterns evoked within this sub-population are involved in differentiating aversive experiences versus avoidance of those aversive experiences. Although this would be in line with theories suggesting that the NAc activity correlates with reward and relief signaling^35, 43, 44^, our results here show that a larger part of the NAc single cell ensembles responds to the aversive events in a positive fashion. Thus, further investigation of the role of specific NAc core single cell sub-populations in processing of social information is required.

Social perception of aversive events (i.e., observational fear^3^) is foundational for empathy-like behavioral phenotypes, which is maladaptive in a number of psychiatric conditions. Indeed, it is well known that individuals with psychiatric pathologies do not find social stimuli as rewarding as neurotypicals do^45^, and studies showed altered neuronal activities in those overlapping regions during social stimuli^5, 7^. Studies in children with ASD showed diminished NAc activity during social cues^45^. Another study in children with ASD evidenced that weak connectivity of NAc contributes to impaired social skills^6^. The NAc belongs to a larger network of overlapping aversive response-social circuitry, with studies showing the involvement of the ventral tegmental area, prefrontal cortex, anterior cingulate cortex and basolateral amygdala in this network^3, 5, 7, 10, 12, 13^. Along with that, human and rodent studies suggest an involvement of the insular cortex in aversive response, empathy-like behaviors and anxiety disorders^46–49^. Therefore, investigations into the structures of these networks play a crucial role in building a better understanding of psychiatric pathologies and providing insights into their treatments.

In conclusion, the present study contributes to the literature of accumbal information processing by showing differential NAc activity patterns to physically and socially experienced aversive events, as well as subpopulations within the NAc core single cell ensembles displaying a bidirectional response to diverse aspects of an aversive stimulus.

## Methods

### Subjects

Male (n=6) and female (n=6) 6-to 8-week-old C57BL/6J mice were obtained from Jackson Laboratories (Bar Harbor, ME; SN: 000664) and housed with five animals per cage. All animals were maintained on a 12h reverse light/dark cycle. Animals received free access to food and water in their homecages. All experiments were conducted in accordance with the guidelines of the Institutional Animal Care and Use Committee (IACUC) at Rowan-Virtua School of Osteopathic Medicine and Vanderbilt University School of Medicine, which approved and supervised all animal protocols.

### Apparatus

Mice were trained and tested daily in individual Med Associates (St. Albans, Vermont) operant conditioning chambers (MED PC operant boxes) fitted with visual stimuli including a standard house light and two cue lights located on each side of the box.

### Behavioral procedures

All “observer” mice were placed in the MED PC operant boxes and received 17 footshocks with a variable inter-stimulus interval (ISI: 15 seconds on average) for the “experienced shock” session. The operant box was divided using a divider and all shocks were administered on the right side of the chamber. The divider was made of see-through plexiglass and holes were drilled to allow the passage of scents. Twenty-four hours after the “experienced shock” session, mice were returned to the same side of the chamber but this time the grid floors were covered with plexiglass and another mouse (performer) was placed in the left side of the chamber. During the “observed shock” session, the performer mouse received 17 footshocks with the same ISI used in the experienced shock session. The mouse that had received the shock in the “experienced shock” session was allowed to observe the performer mouse. The performer mouse was the same sex of the observer mouse. All shocks were set to 1.0 mA in intensity and 0.5 seconds in duration.

### Surgical procedure

Ketoprofen (5mg/kg; subcutaneous injection) was administered at least 30 minutes before surgery. Under Isoflurane anesthesia, mice were positioned in a stereotaxic frame (Kopf Instruments) and the NAc core (bregma coordinates: anterior/posterior, +1.4 mm; medial/lateral, +1.0 mm; dorsal/ventral, −3.8 mm; 0° angle) was targeted. Ophthalmic ointment was applied to the eyes. Using aseptic technique, a midline incision was made down the scalp and a craniotomy was made using a dental drill. A 10 mL Nanofil Hamilton syringe (WPI) with a 34-gauge beveled metal needle was used to infuse viral constructs. A calcium indicator virus, GCamP (AAV1.Camk2a.GCaMP6m.WPRE.SV40, Inscopix), was infused at a rate of 50 nL/min for a total of 500 nL. Following infusion, the needle was kept at the injection site for seven minutes and then slowly withdrawn. After the virus injection, using a 27-gauge needle (0.4 mm in diameter), a pathway for the subsequent lens implantation was opened. Permanent implantable Proview integrated lenses (baseplates with attached GRIN Lenses 0.6mm diameter, 7.3mm length, Inscopix) were implanted in the NAc. Lenses were positioned above the viral injection site (bregma coordinates: anterior/posterior, + 1.4 mm; medial/lateral, + 1.0 mm; dorsal/ventral, −3.7 mm; 0° angle) and were cemented to the skull using a C&B Metabond adhesive cement system. Animals were allowed to recover for a minimum of six weeks to ensure efficient viral expression before commencing experiments.

### Histology

Subjects were deeply anesthetized with an intraperitoneal injection of Ketamine/Xylazine (100mg/kg/10mg/kg) and transcardially perfused with 10 mL of PBS solution followed by 10 mL of cold 4% PFA in 1x PBS. Animals were quickly decapitated; the brain was extracted and placed in 4% PFA solution and stored at 4 °C for at least 48 hours. The brains were then transferred to a 30% sucrose solution in 1x PBS and allowed to sit until the brains sank to the bottom of the conical tube at 4::J°C. After sinking, brains were sectioned at 35μm on a freezing sliding microtome (Leica SM2010R). Sections were stored in a cryoprotectant solution (7.5% sucrose + 15% ethylene glycol in 0.1 M PB) at -20::J°C until immunohistochemical processing. Sections were incubated for 5::Jminutes with DAPI (NucBlue, Invitrogen) to achieve counterstaining of nuclei before mounting in Prolong Gold (Invitrogen). Following staining, sections were mounted on glass microscope slides with Prolong Gold antifade reagent. Fluorescent images were taken using a Keyence BZ-X700 inverted fluorescence microscope (Keyence), under a dry 10x objective (Nikon). The injection site location and the fiber implant placements were determined via serial imaging in all animals.

### Single-photon calcium imaging via miniscopes

For all single cell imaging, we used endoscopic miniature scopes (nVista miniature microscope, Inscopix) combined with a calcium indicator, GCaMP6m (GCamP (AAV1.Camk2a.GCaMP6m.WPRE.SV40, Inscopix), in order to record single cell activity in the NAc core *in vivo*. During each behavioral session, the miniscope was attached to the integrated lens baseplates implanted previously. The imaging parameters (gain, LED power, focus) were determined for each animal to ensure recording quality and were kept constant throughout the study. The imaging videos were recorded at 10 frames per second (fps). At the end of the recording session, the miniscope was removed and the baseplate cover was replaced. During each session, important events such as stimulus or outcome presentation times were recorded via transistor-transistor-logic (TTL) signals sent from the MedPC behavioral box to the Inscopix data acquiring computer.

### Single-photon calcium imaging analysis

Data was acquired at 10 frames per second using an nVista miniature microscope (Inscopix). TTLs from MedPC were directly fed to the nVista system, which allowed alignment to behavioral timestamps without further processing. The recordings were spatially down-sampled by a factor of 2 and corrected for motion artifacts using the Inscopix Data Processing Software (IDPS v1.3.1). The ΔF/F values were computed for the whole field view as the output pixel value represented a relative percent change from the baseline. We used Nonnegative Matrix Factorization - Extended (CNMF-e^50, 51^) to identify and extract calcium traces from individual cells (CNMF-e cell detection parameters: patch_dims = 50, 50; K = 20; gSiz = 20; gSig = 12; min_pnr = 20; min_corr = 0.8; max_tau = 0.400). Raw CNMFe traces were used for all analyses. The spatial mask and calcium time series of each cell were manually inspected using the IDPS interface. Cells found to be duplicated or misdetected due to neuropils or other artifacts were discarded. The raw ΔF/F data were exported and used for TTL analysis, in which we cropped the data around each significant event (cue presentations; TTL) and z-scored it in order to normalize for baseline differences. Z-scores were calculated by taking the pre-TTL ΔF/F values as a baseline (z-score = (TTLsignal - b_mean)/b_stdev, where the TTL signal is the ΔF/F value for each post-TTL time point, b_mean is the baseline mean, and b_stdev is the baseline standard deviation). Using the z-scored traces, we then calculated whether the cell response to the experienced and observed footshock stimuli was significant in order to determine responsive and non-responsive cells as well as the direction of the response (positive or negative). For this analysis, we calculated averaged area under the curve (AUC) shock cell responses during a 2-second post-TTL window. We then ran one-tailed independent t-tests to determine that the response was significantly different than the baseline.

### Longitudinal registration

We used the longitudinal registration pipeline, defined in the Inscopix Data Process Software (IDPS) Guide, to identify the same cell across recording sessions in longitudinal series. Cell sets are preprocessed to generate a cell map which is then aligned to the first cell map (the reference). The images of the first cell set are defined as the global cell set against which the other cell sets are matched. We then find the pair of cell images between the global cell set and other aligned cell sets that maximizes the normalized cross correlation (NCC). The program then generates an output that aligns the same cell from across sessions.

### Hierarchical clustering

Using the calcium traces from all the cells detected in the “experienced shock” versus “observed shock” sessions, we employed a hierarchical clustering approach to group cells based on their shock responses in an unbiased way. We used the “clustergram” Matlab function and the “correlation” distance metric to group the cell activity. We then combined all the cells that were clustered in Node 1 and Node 2 to visualize the group characteristics (e.g., positive response vs. negative response to shock)

## Acknowledgements

This work was supported by NIH grants R21MH132052 and KL2TR002245 to M.G.K.

## Author Contributions

MGK conceptualized the study. MGK and JEZ performed the viral surgeries and ran the behavioral and imaging experiments. MGK and OD analyzed the data. MGK and OD wrote the manuscript. All authors edited and approved the final version of the manuscript.

## Data availability statement

All data in the manuscript or the supplementary material are available from the corresponding author upon reasonable request. Correspondence and requests for materials should be addressed to Munir Gunes Kutlu

**Supp Figure 1.**
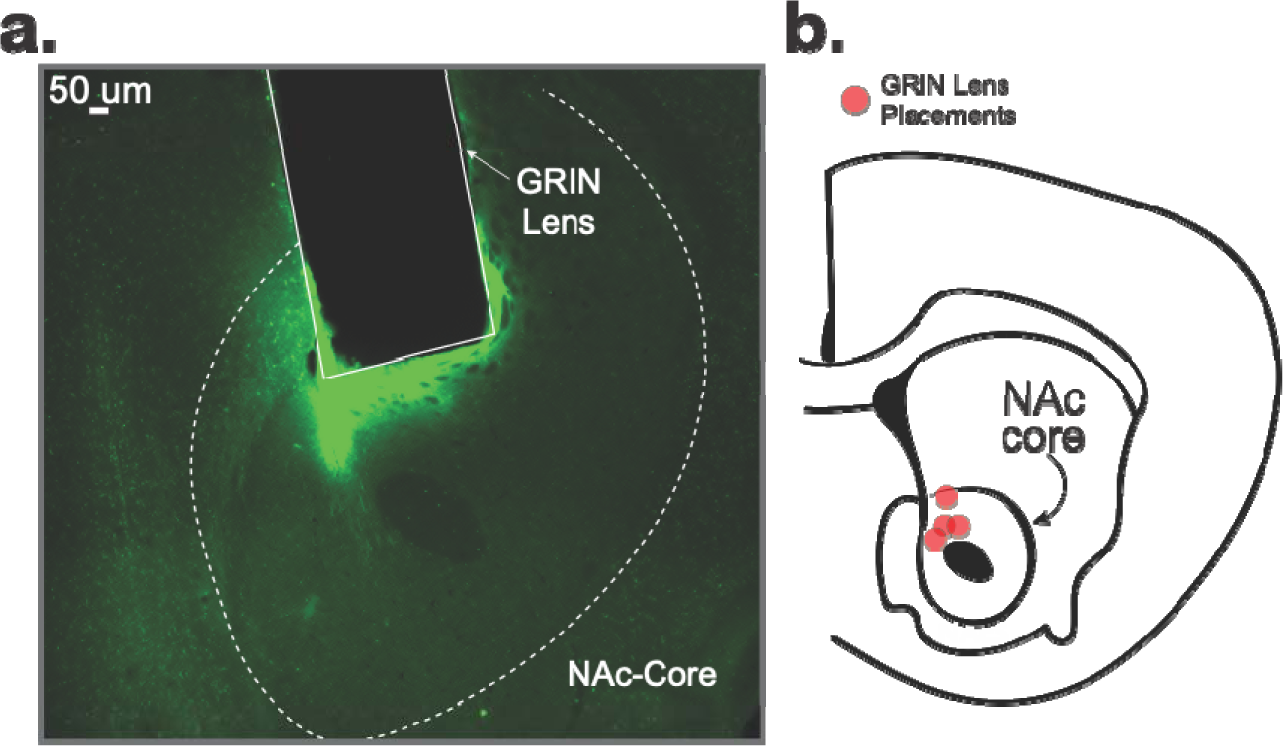
Representative histology and single-cell calcium traces. **(a)** Representative image showing the expression of GCaMP in the NAc core and the placement of the GRIN lens relative to the viral expression. **(b)** Map showing GRIN lens placement in all mice.

**Supp Figure 2.**
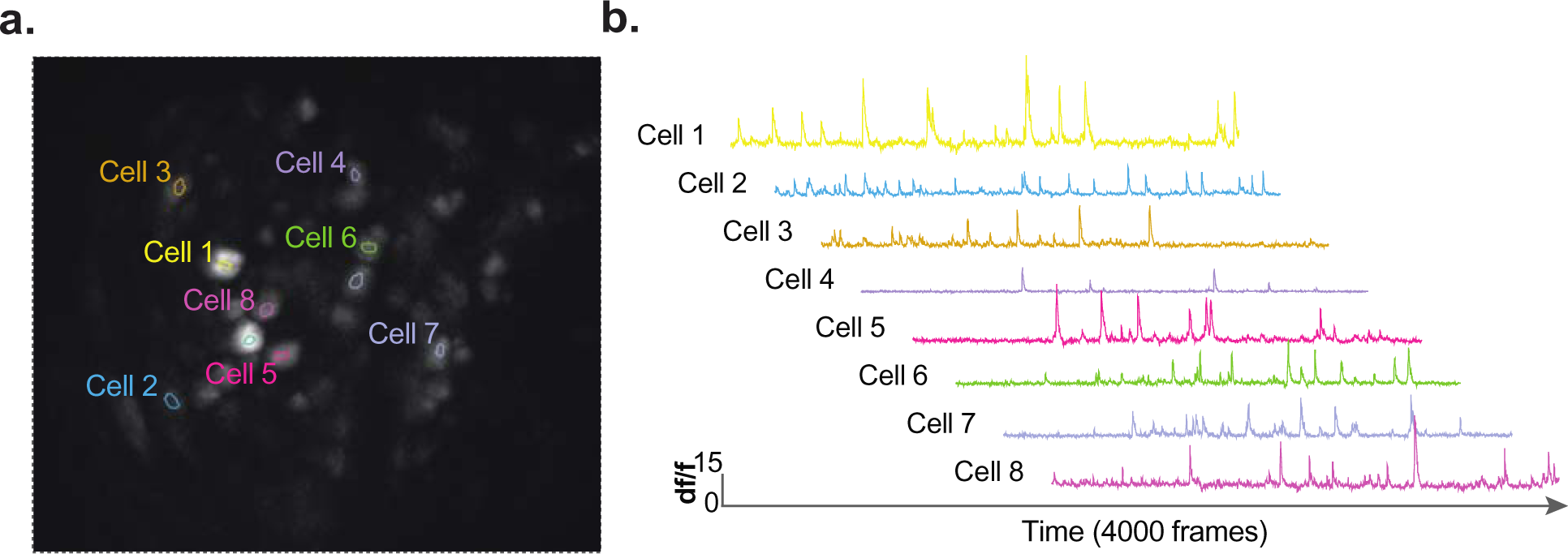
**(a)** Representative cell map showing detected single cells from an individual animal. **(b)** Representative calcium traces from individual neurons identified in one animal.

**Supp Figure 3.**
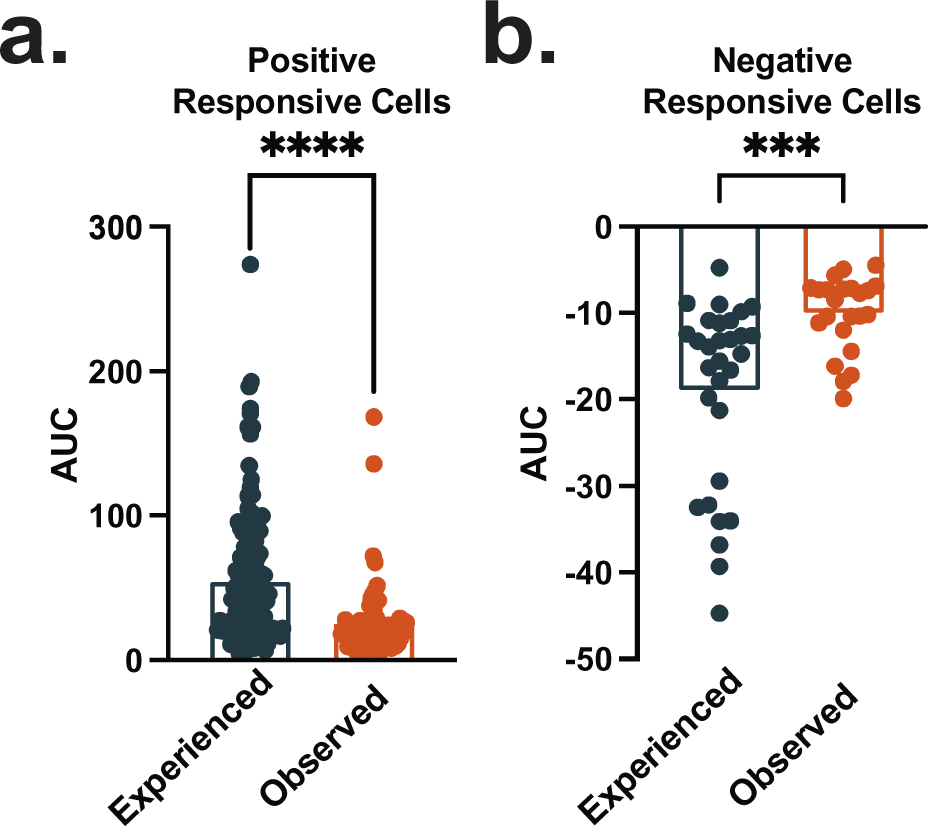
Single cell responses to the experienced footshocks were stronger compared to the responses to observed footshocks. (**a**) Single cell responses from the cells that showed a significant positive response were significantly stronger during the experienced footshock session (unpaired t-test, *t*_209_=5.037, *p*<0.0001, n=71-140 cells). (**b**) Single cell responses from the cells that showed a significant negative response were significantly stronger during the experienced footshock session (unpaired t-test, *t*_51_=3.751, *p*=0.0005, n=23-30 cells). Data represented as mean ± S.E.M. *** *p* < 0.001, **** *p* < 0.0001.

**Supp Figure 4.**
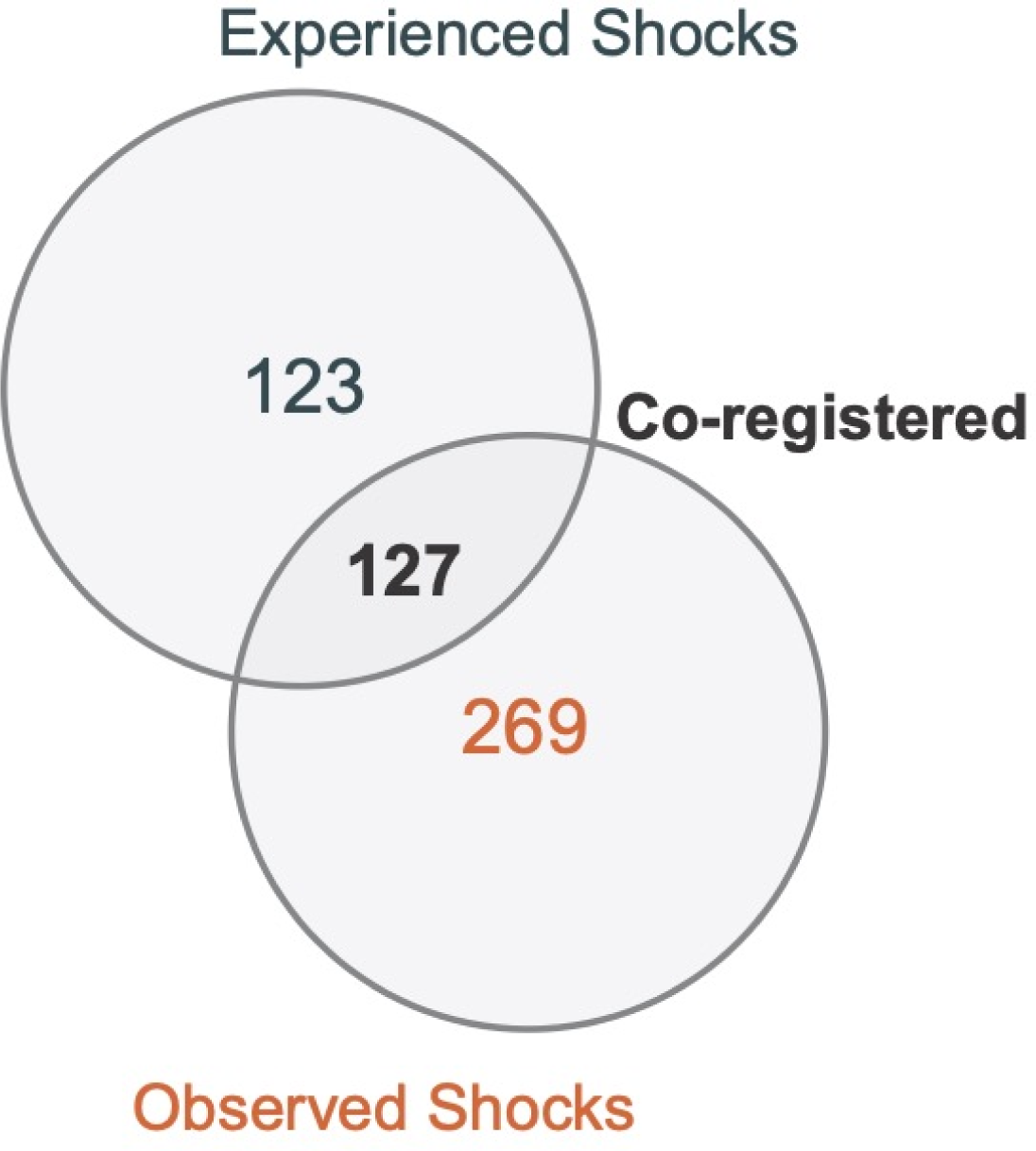
Experienced and observed footshock responsive cell ensembles showed a large overlap. Of all cells that detected during the experienced versus observed footshock sessions, there was a large number of cells that were detected in both sessions (n = 127 cells).

**Supp Figure 5.**
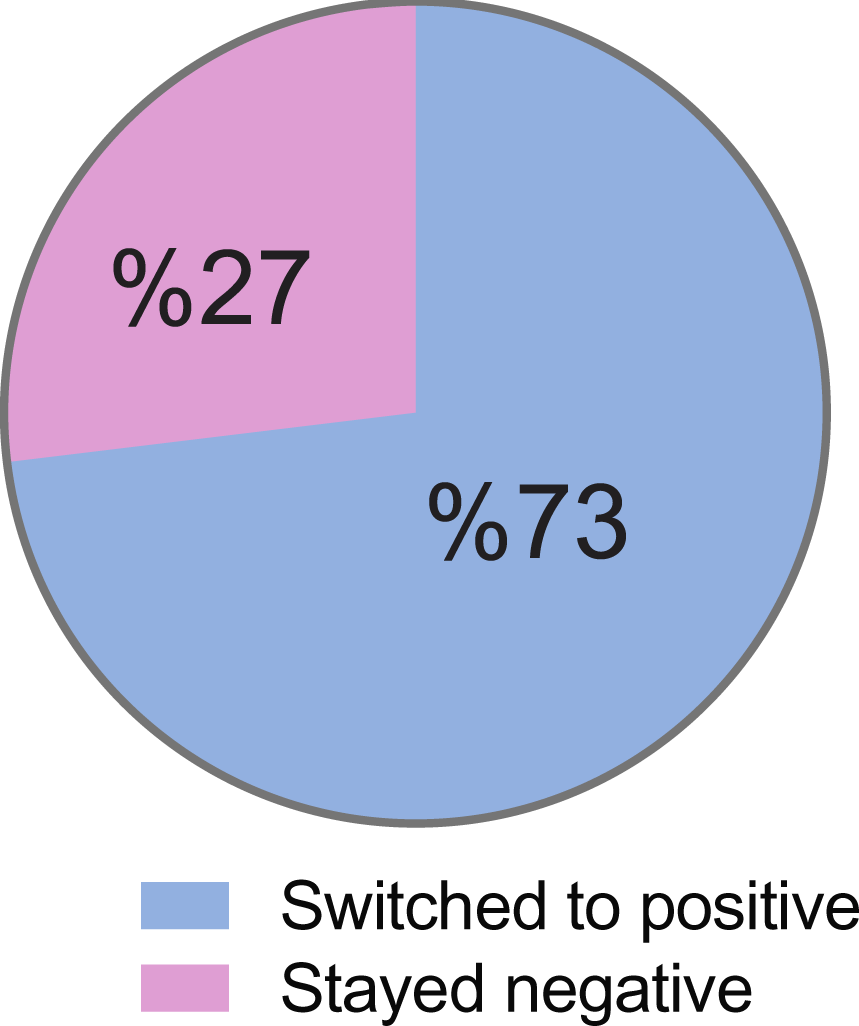
Majority of the cells that showed a negative response to the experienced footshocks switched to a positive response when the mice observed another mouse receiving footshocks (73%).

## Notes

### Competing Interest Statement

The authors have declared no competing interest.

